# Why structural divergence varies among residues in enzyme evolution: contributions of mutation, stability, and activity constraints

**DOI:** 10.1101/2025.10.07.680993

**Authors:** Julian Echave, Mathilde Carpentier

**Affiliations:** Instituto de Ciencias Físicas (ICIFI-CONICET), Universidad Nacional de San Martín Martín de Irigoyen 3100, 1650 San Martín, Buenos Aires, Argentina; Institut de Systématique, Évolution, Biodiversité (ISYEB, UMR 7205) CNRS-MNHN-SU-EPHE-UA, Sorbonne Université 57 rue Cuvier, 75005 Paris, France

## Abstract

In enzyme evolution, structural divergence varies among residues, forming residue-dependent structural divergence profiles. The evolutionary constraints that determine these profiles remain unclear. We build on the Mutation-Stability-Activity (MSA) model, a mechanistic mutation-selection model previously developed for sequence evolution. In the MSA model, mutations become fixed or are lost depending on their effects on stability and activity, with parameters *a*_*S*_ and *a*_*A*_ controlling selection on stability and activity, respectively. The Linearly Forced Elastic Network Model (LFENM) is used to calculate mutational effects on structure, stability, and activity. As substitutions accumulate, structural changes build up unevenly across residues, producing a structural divergence profile that depends on how each residue responds to mutation and on how strongly selection acts on stability and activity. Applied to 34 enzyme families, the MSA model recapitulates observed structural divergence profiles, and nested model comparisons show that mutation, stability, and activity constraints each contribute. However, the balance among these constraints varies widely across families: mutation always contributes substantially, but stability and activity contributions range from negligible to dominant, so any of the three can prevail in a given family. These variations have distinct origins: the mutation contribution depends on how unevenly the protein’s flexibility is distributed across residues, while the stability and activity contributions depend on how strongly selection acts, as quantified by *a*_*S*_ and *a*_*A*_. The MSA model thus recovers family-specific selection strengths from structural divergence profiles, suggesting these profiles encode information not only about enzyme architecture but also about the selective regime under which enzymes evolve.

## 1 Introduction

Over the past two decades, biophysical models have provided mechanistic explanations for a range of sequence evolution patterns, including variation of substitution rates among sites, residue-dependent substitution matrices, and covariation (Echave and Wilke, 2017) (for recent developments, see also (Ferreiro et al., 2024; André, 2025) and references therein). Much work remains, but a substantial theoretical foundation exists. For structural evolution, the picture is very different: some empirical work has characterized patterns protein structure divergence (Chothia and Lesk, 1986; Wood and Pearson, 1999; Leo-Macias et al., 2005; Illergård et al., 2009; Friedland et al., 2009; Mahajan et al., 2014; Vetrivel et al., 2019; Echave and Carpentier, 2025), but biophysical models that explain the observed patterns are scarce.

In particular, it is not understood why structural divergence varies among residues. Some positions change substantially across homologues while others remain nearly fixed, producing residue-dependent structural divergence profiles. Our recent empirical analysis of 34 families of functionally conserved homologous enzymes characterized these profiles in detail (Echave and Carpentier, 2025): structural divergence increases with backbone flexibility and with distance from catalytic residues. Decomposing the profiles into flexibility-dependent and distance-dependent components revealed that both non-functional and functional constraints influence them. However, this empirical analysis falls short of a mechanistic understanding of what these constraints are and how they operate.

Diffusion models of structural change have been developed (Gutin and Badretdinov, 1994; Grishin, 1997; Challis and Schmidler, 2012; Golden et al., 2017; Larson et al., 2020). These models are useful for estimating evolutionary distances and trees, but they assume that all residues evolve by the same process, so that they cannot account for the variation of structural divergence among residues.

Toy models of protein evolution that incorporate mutation, stability, and ligand-binding constraints have also been developed (Nelson and Grishin, 2014, 2016). Because these models simulate the full folding process, they can in principle predict that structural divergence varies among residues. However, being based on toy proteins, their predictions cannot be compared quantitatively with observed divergence profiles of real protein families.

Another approach to structural divergence is based on the Linearly Forced Elastic Network Model (LFENM), which models the structural effect of mutations on a known protein structure (Echave, 2008). While limited to small structural changes, this approach can be applied to real proteins and compared directly with observed divergence profiles. Using LFENM, we showed that the shape of structural divergence profiles is well explained by the mechanical response of the protein to random mutations, without invoking selection (Echave and Fernández, 2010); this is essentially a mutation-only model (MM). Subsequently, incorporating selection on stability revealed that stability constraints also contribute to structural divergence, amplifying the profile without changing its shape (Marcos and Echave, 2020); this is a mutation-stability model (MS).

If mutation and stability constraints were sufficient to explain structural divergence profiles, we would not expect the influence of functional constraints found in our empirical analysis (Echave and Carpentier, 2025). Moreover, in previous work we developed a Mutation-Stability-Activity model (MSA) for enzyme sequence evolution and showed that activity constraints are needed to fully account for sequence divergence patterns (Echave, 2019, 2021), suggesting that they may also be needed to explain structural divergence patterns. Here, we extend the MSA model to structural evolution to predict residue-dependent structural divergence profiles. By decomposing the predicted profiles into mutation, stability, and activity components, we can identify which constraints operate in each enzyme family, quantify their relative importance, and trace their variation among families to underlying biophysical and selective determinants. (Note that throughout this work, MSA denotes the Mutation-Stability-Activity model, not multiple sequence alignment.)

## 2 Results

We use the MSA model to study the roles of mutation, stability, and activity constraints in residue-dependent structural divergence profiles. After an overview of the model, we show that MSA, with only two free parameters, captures observed profiles as accurately as a flexible empirical model; that all three constraints contribute to the profiles; that their relative contributions vary widely among enzyme families; and that this variation is explained by flexibility heterogeneity for the mutation contribution and by selection strengths for the stability and activity contributions.

### 2.1 Overview of the Mutation-Stability-Activity model

We study 34 families of functionally conserved enzymes from the M-CSA database, previously curated in (Echave and Carpentier, 2025). Each family consists of a reference enzyme, for which the catalytic mechanism and active-site residues have been experimentally determined, and a set of homologous structures. Superimposing the homologous structures onto the reference and measuring the per-residue C*α* RMSD yields a structural divergence profile, a residue-by-residue map of how much each position has shifted over evolutionary time. The MSA model aims to predict these profiles from first principles.

The MSA model represents evolution as a mutation-selection process. The process starts at an ancestral wild-type protein, which we take here to be the M-CSA reference for the studied family. Then, at each evolutionary time step, a random mutation is introduced at a randomly chosen residue, displacing the structure from its wild-type conformation by Δ**r**^0^, changing the folding free energy by ΔΔ*G*, and changing the catalytic activation energy by ΔΔ*G*^‡^. The mutation either fixes or is lost according to the fixation probability (Eq. 2 in Methods):

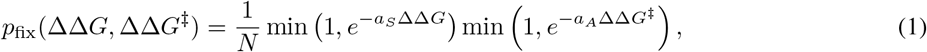

where *N* is the population size, and *a*_*S*_ and *a*_*A*_ are parameters that control the strength of selection against destabilizing and deactivating mutations, respectively. Selection is purely negative: mutations that do not compromise stability or activity (ΔΔ*G* ≤ 0 or ΔΔ*G*^‡^ ≤ 0) fix at the neutral rate 1*/N*, while deleterious mutations fix at rates that decrease exponentially with the magnitude of the effect and the corresponding selection parameter. As substitutions accumulate, structural changes build up unevenly across residues, producing the structural divergence profile.

We calculate Δ**r**^0^, ΔΔG, and ΔΔG^‡^ using the Linearly Forced Elastic Network Model (LFENM) (Echave, 2008; Echave and Fernández, 2010). LFENM represents the reference protein as a network of C*α* beads connected by harmonic springs. A mutation is modelled by adding random perturbations to the lengths of the springs that connect the mutated site with its neighbours. Relaxation of the mutant to a new equilibrium displaces the structure by Δ**r**^0^. The mutation also creates local stress in the network; the structure partially relaxes, but some stress remains, and this residual stress is ΔΔG. Finally, because the structural change shifts the active-site geometry away from the wild-type conformation, and assuming that the wild-type conformation is optimal for catalysis (preorganisation), the mutant must distort its active site back to the wild-type geometry to achieve catalysis; the energetic cost of this distortion is ΔΔG^*‡*^. Full derivations are given in Methods (Eqs. 6–8), and a minimal illustration is given in Supplementary Section S1 and Supplementary Figure S1.

In principle, predicting a structural divergence profile requires simulating the accumulation of many substitutions. However, because the RMSD of all residues scales by the same factor as substitutions accumulate, the shape of the profile does not change with the number of substitutions, and the single-substitution ensemble is sufficient. To generate this ensemble, we introduce multiple random mutations at each residue position using LFENM. Each mutant has an associated structural displacement, and its probability in the ensemble is proportional to its fixation probability. The predicted per-residue RMSD is the root mean square displacement over this ensemble. The only free parameters are a_S_ and a_A_, which we estimate for each enzyme family by Bayesian inference, fitting predicted to observed structural divergence profiles. Full details are in Methods (Section 4).

### 2.2 The MSA model captures empirical patterns mechanistically

We first assess how well the MSA model captures observed structural divergence profiles, comparing its predictions with observations and with our previous empirical model M12 (Echave and Carpentier, 2025), a shape-constrained additive model that fits observed divergence as a function of residue flexibility and distance to the active site.

For each of the 34 enzyme families of this study, we quantified observed structural divergence by the per-residue C*α* RMSD across homologues, and predicted RMSD profiles using the MSA model as described in Section 2.1. The predicted profile for each family was obtained by first generating a predicted RMSD profile for each draw from the posterior distribution of (a_S_, a_A_), estimated by Bayesian inference, and then averaging these profiles across draws. To focus on how divergence varies among residues rather than its overall magnitude, both observed and predicted RMSD profiles were log-transformed and centred by subtracting the protein mean, yielding nlRMSD_obs_ and nlRMSD_MSA_ profiles.

The MSA model captures observed structural divergence as accurately as M12, and the two models’ predictions are very similar. The average correlation between predicted and observed profiles is 0.66 ± 0.02 for the MSA model and 0.69 ± 0.02 for M12 (ranges 0.43 − 0.83 and 0.45 − 0.86, respectively; Fig. 1A and Supplementary Table S3). The average correlation between MSA and M12 predictions is 0.94 ± 0.01 (range 0.8 − 0.99; Fig. 1B).

**Figure 1:**
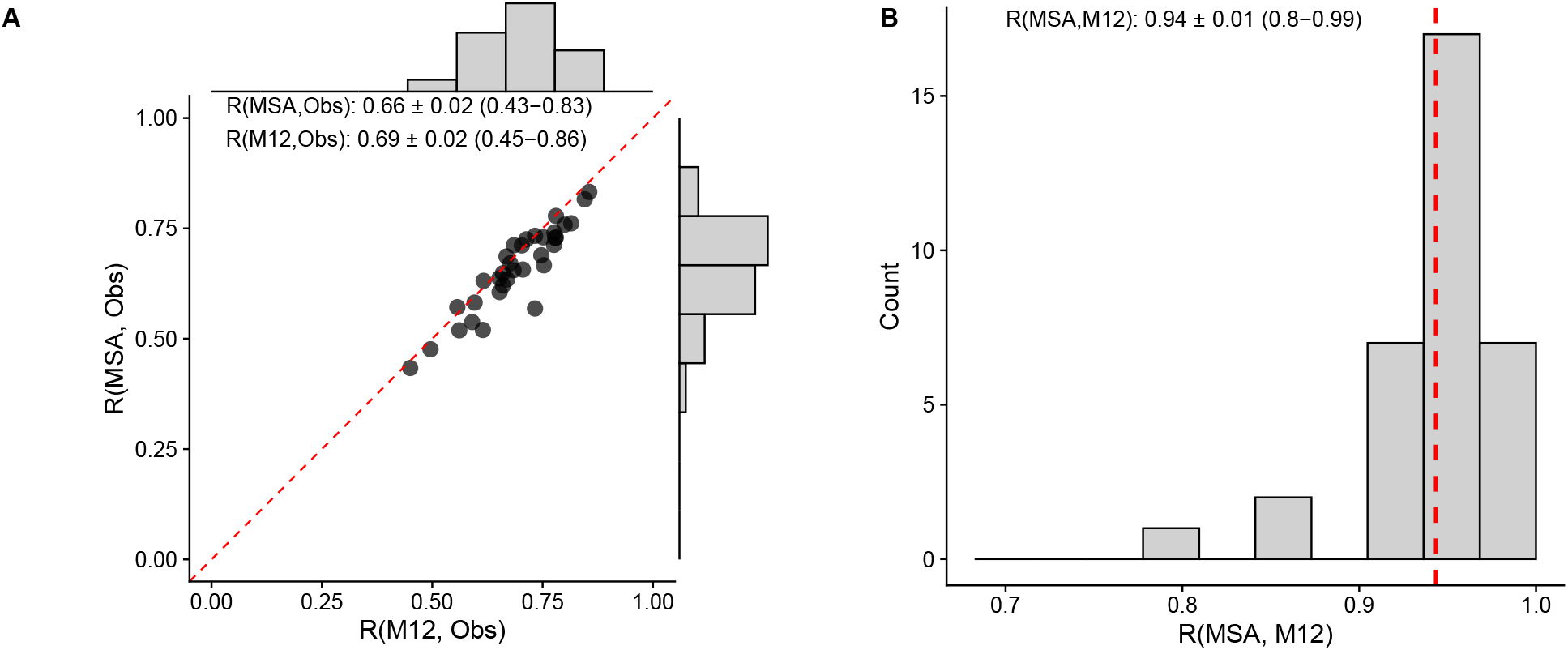
Comparison of MSA and empirical model performance. (A) Scatter plot comparing correlations between model predictions and observed structural divergence profiles, measured as centred log–RMSD (nlRMSD), across 34 enzyme families. The x-axis shows correlations between the empirical model (M12) and observed profiles, the y-axis shows correlations between MSA predictions and observed profiles. Each point represents one enzyme family The dashed red line indicates perfect agreement. Marginal histograms show the distribution of prediction–observation correlations for each model. Annotations show mean ± SEM and range (min–max) for each model’s correlations with observations. (B) Histogram showing the distribution of correlations between MSA and M12 predictions across all enzyme families. The red dashed line indicates the mean correlation. Annotation shows mean ± SEM and range (min–max).

The empirical model M12 describes how structural divergence depends on flexibility and distance to the active site and attributes this to non-functional and functional constraints, but without a mechanistic model to support this interpretation. The close agreement between MSA and M12 profiles indicates that the MSA model captures the same dependencies mechanistically. The following sections show how mutation, stability, and activity constraints contribute to structural divergence profiles.

### 2.3 Three constraints influence structural divergence profiles

To determine whether all three constraints influence structural divergence profiles, we compared four models of increasing complexity: a baseline of uniform divergence (M0), a mutation model (MM) that allows divergence to vary among residues due to differences in structural response to mutation but includes no selection, a mutation-stability model (MS) that adds selection against destabilizing mutations, and the full MSA model that further adds selection against deactivating mutations. If each step from M0 to MM to MS to MSA improves the fit, it means that the added constraint leaves a detectable trace in the observed profiles.

Computationally, the nested models are special cases of MSA: MM corresponds to setting a_S_ = a_A_ = 0, MS to setting a_A_ = 0, and the full MSA retains both selection parameters. We quantify fit using explained deviance *D*^2^, the fraction of variance in observed nlRMSD profiles explained by the model (*D*^2^ = 0 for the baseline, *D*^2^ = 1 for a perfect fit; *D*^2^ reduces to *R*^2^ for linear models; see Methods).

To illustrate the nested comparison with concrete examples before turning to aggregate results, we highlight three enzyme families (Fig. 2A). For the aldo/keto reductase family (M-CSA ID 858), *D*^2^ increases from 0.09 (MM) to 0.56 (MS) with no further improvement in MSA (0.56), indicating that mutation and stability constraints contribute to the profile but activity constraints do not. For the metallo-beta-lactamase family (M-CSA ID 15), *D*^2^ increases at each step (MM: 0.23, MS: 0.40, MSA: 0.59), indicating that all three constraints contribute. For the ribonuclease U2 family (M-CSA ID 908), *D*^2^ increases from 0.11 (MM) to 0.17 (MS) to 0.65 (MSA), indicating that mutation and activity constraints contribute while stability constraints have a marginal effect. Although not all constraints contribute detectably in every family, these three cases illustrate that each of the three constraints can contribute to structural divergence profiles.

**Figure 2:**
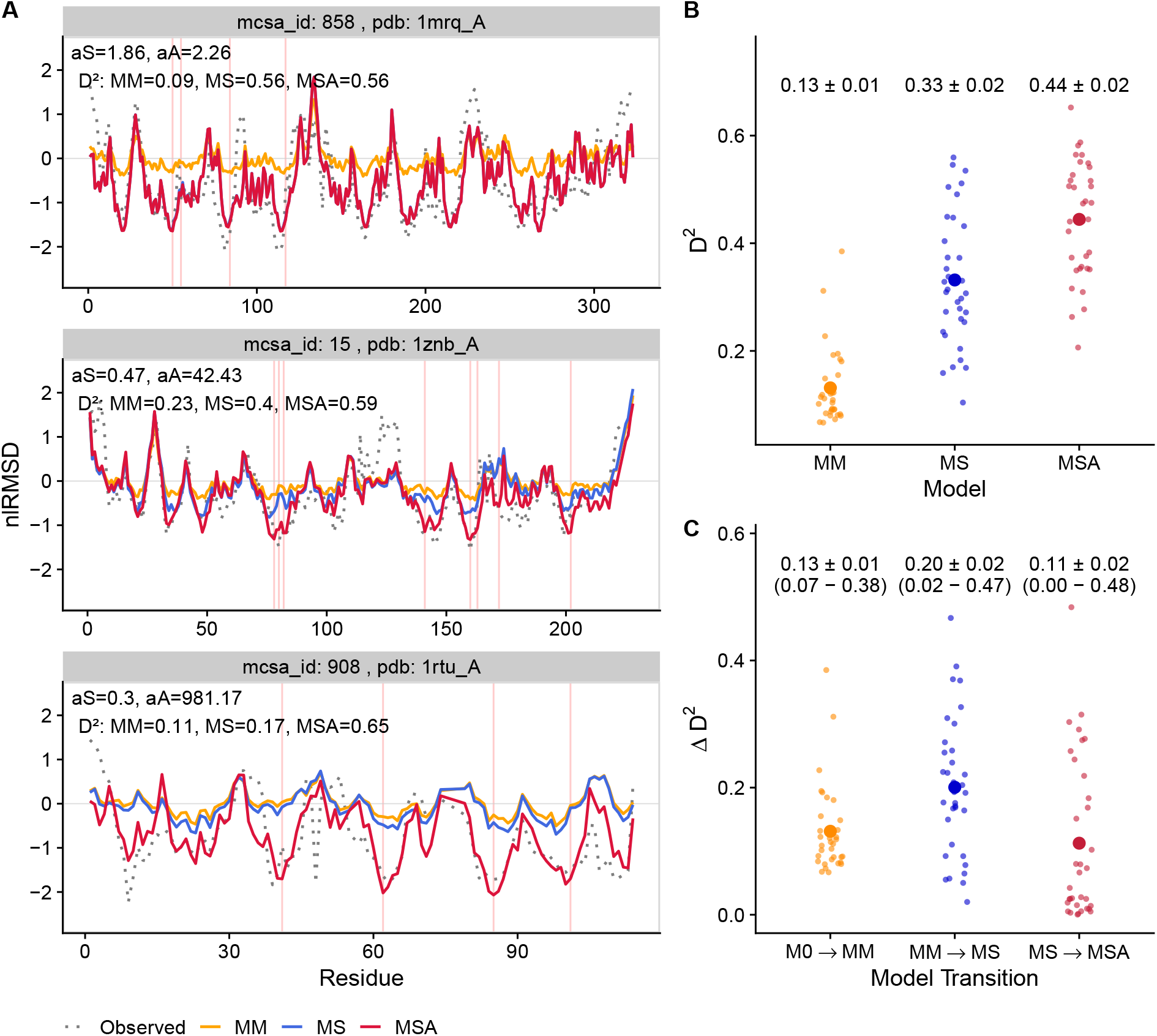
Nested model comparison of structural divergence profiles across enzyme families. (A) Residue-dependent structural divergence profiles for three enzyme families. Each panel shows observed centred log–RMSD (nlRMSD, gray dashed line) and predictions from M0 (residue-independent uniform divergence, gray line), MM (mutation-only model, orange), MS (mutation–stability model, blue), and MSA (mutation–stability–activity model, red). Vertical red lines mark active site residues. Annotations show posterior mean estimates of a_S_ and a_A_ from MSA, together with the observed deviance explained by each model *D*^2^. (B) Distribution of deviance explained by models MM, MS, and MSA across all enzyme families. (The deviance explained by M0 is, by definition, zero, and is not shown.) Small points represent individual enzyme families; large points indicate means over families. Means ± standard errors are shown above each distribution. (C) Distribution of incremental explained deviance from sequentially adding mutation (M0 → MM), stability (MM → MS), and activity (MS → MSA) constraints. Small points represent individual enzyme families; large points indicate means over families. Means ± standard errors and ranges (min–max) are shown above each distribution.

Across the full dataset, each constraint significantly improves fit on average (Fig. 2B,C). Mean *D*^2^ values across all 34 families increase from 0.13 ± 0.01 (MM) to 0.33 ± 0.02 (MS) to 0.44 ± 0.02 (MSA), with each mean increment significant at *p <* 0.001. Thus, on average, all three constraints contribute to structural divergence profiles.

However, not all constraints contribute detectably in every family. While the mean increments are all significant, the Δ*D*^2^ from adding any given constraint varies widely across families (Fig. 2C). Mutation always contributes (Δ*D*^2^ range 0.07–0.38), but the increments from adding stability or activity constraints can be near zero, indicating that these constraints may have negligible effects in particular families.

These results establish that the three MSA constraints influence structural divergence profiles. The improvement from M0 to MM demonstrates that mutational effects create residue-dependent variation beyond uniform divergence, so structural divergence would be residue-dependent even in the absence of selection. The improvements from MM to MS and from MS to MSA demonstrate that both stability and activity constraints also contribute to the observed profiles. Since which constraints contribute varies among families, all three need to be considered when analysing structural evolution in any given enzyme family.

The nested comparison establishes whether each constraint contributes, but does not quantify how much each one contributes to the profile. To quantify individual contributions, we turn to a decomposition approach.

### 2.4 Constraint contributions vary among enzymes

To quantify the individual contributions of mutation, stability, and activity constraints to structural divergence profiles, we decompose the predicted profile into three component profiles, *ϕ*_M_, *ϕ*_S_, and *ϕ*_A_, such that nlRMSD_MSA_ = *ϕ*_M_ + *ϕ*_S_ + *ϕ*_A_ (see Methods for definitions).

We measure how much a component profile contributes to the divergence profile by its standard deviation across residues, giving three values per family, SD(*ϕ*_M_), SD(*ϕ*_S_), and SD(*ϕ*_A_). Dividing these three values by their sum gives the relative contributions, which quantify the balance among constraints for a family.

In our three example families, mutation constraints contribute substantially in all cases, but the contributions of stability and activity constraints vary widely (Fig. 3A). In the aldo/keto reductase family (M-CSA ID 858), mutation contributes 0.39, stability contributes most of the rest (0.60), and activity is marginal (0.01). In the metallo-beta-lactamase family (M-CSA ID 15), mutation again contributes substantially (0.48), with stability (0.19) and activity (0.33) sharing the remainder. In the ribonuclease U2 family (M-CSA ID 908), mutation contributes 0.34, stability is small (0.09), and activity contributes most of the rest (0.57).

**Figure 3:**
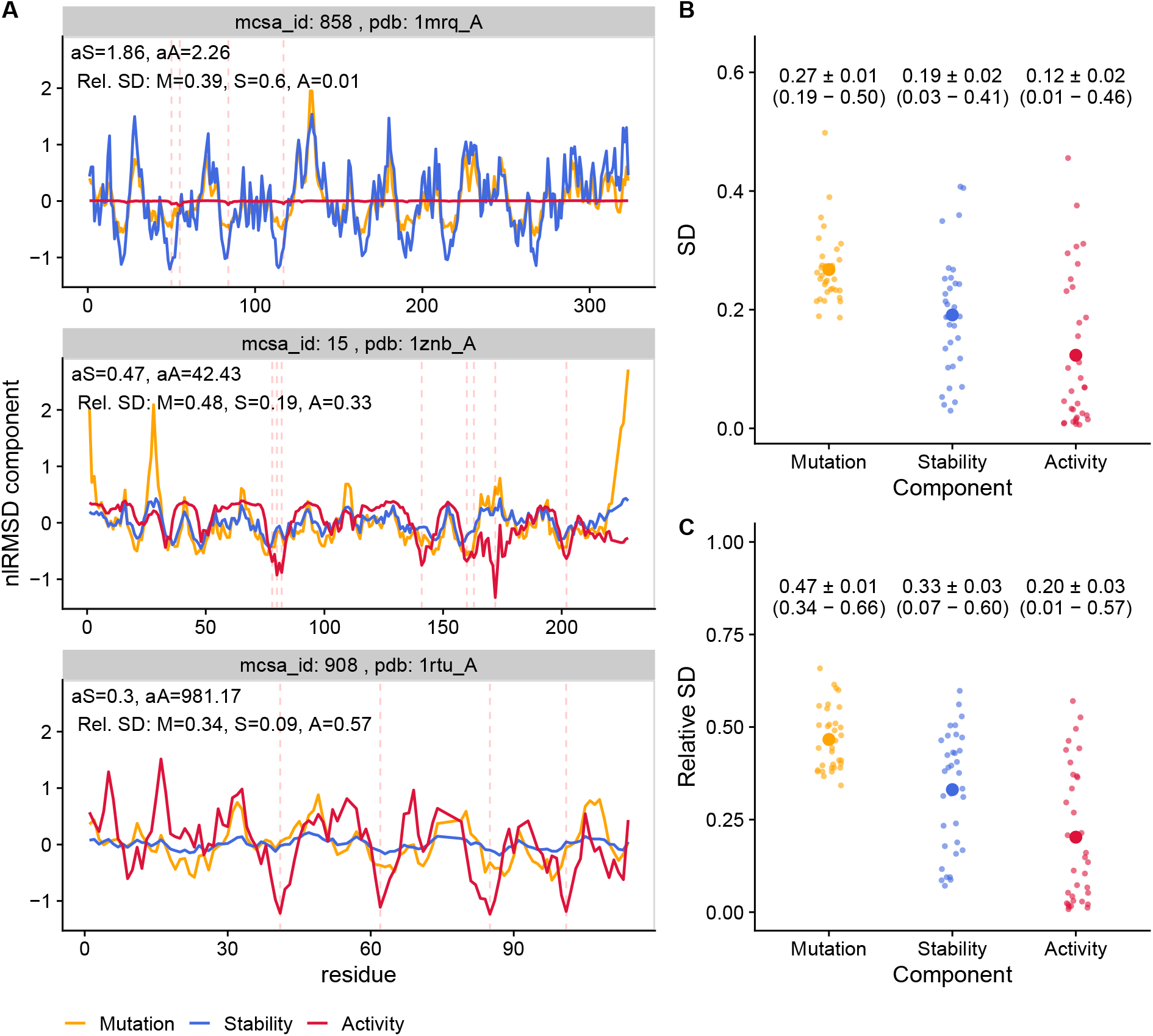
Decomposition of divergence profiles into mutation, stability, and activity components. MSA profiles are decomposed into a sum of three components (nlRMSD = *ϕ*_M_ + *ϕ*_S_ + *ϕ*_A_). Each component’s contribution to the profile is quantified by its standard deviation across residues: SD(*ϕ*_M_) for mutation, SD(*ϕ*_S_) for stability, and SD(*ϕ*_A_) for activity. The relative contribution is defined as the component’s SD divided by the sum of the SDs of all components. (A) Component profiles for three enzyme families showing mutation (orange), stability (blue), and activity (red) contributions. Vertical dashed red lines mark active site residues. Annotations show posterior mean estimates of a_S_ and a_A_, together with the relative SD of each component. (B) Distributions of component contributions (SD) across the dataset. (C) Distributions of relative component contributions (relative SD) across the dataset. In Panels B and C, small points represent individual families; large points indicate means over families. Means ± SEM and ranges (min–max) are shown above the distributions.

Across the full dataset, mutation constraints contribute most on average, accounting for about half the profile (relative SD: 0.47 ± 0.01), followed by stability (0.33 ± 0.03) and activity (0.20 ± 0.03) constraints (Fig. 3B,C and Supplementary Table S3). Beyond these averages, the components vary differently among families. Mutation contributions are consistently substantial (relative contribution 0.34–0.66), whereas stability and activity contributions vary widely (0.07–0.60 and 0.01–0.57, respectively), so that either can range from marginal to dominant.

There is thus no universal hierarchy of constraints: while mutation constraints always contribute substantially, the roles of stability and activity constraints are family-specific, and any of the three can become the prevailing constraint in a given family. What determines these different balances is the subject of the following section.

### 2.5 Constraint balance is determined by protein architecture and selection strengths

Having established that the balance among constraints varies widely across enzyme families, we now ask what determines this variation.

Theoretically, in the MSA model, the RMSD profile is determined by the distributions of the mutational effects Δ**r**^0^, ΔΔ*G*, and ΔΔ*G*^*‡*^, and by the selection parameters a_S_ and a_A_. Variation among enzyme families in the mutation, stability, and activity contributions can therefore arise from differences in the distributions of mutational effects, or in the selection parameters, or both. Since in LFENM the distributions of mutational effects are determined by protein architecture (structure and active-site location), variation among families can arise only from differences in protein architecture or in selection strengths.

Empirically, each contribution is primarily determined by a single factor (Fig. 4). The mutation contribution correlates strongly with flexibility heterogeneity, SD(lRMSF) (Spearman *ρ* = 0.99; Fig. 4A). The stability contribution correlates strongly with the stability selection parameter a_S_ (*ρ* = 0.93; Fig. 4B). The activity contribution correlates strongly with the activity selection parameter a_A_ (*ρ* = 0.94; Fig. 4C).

**Figure 4:**
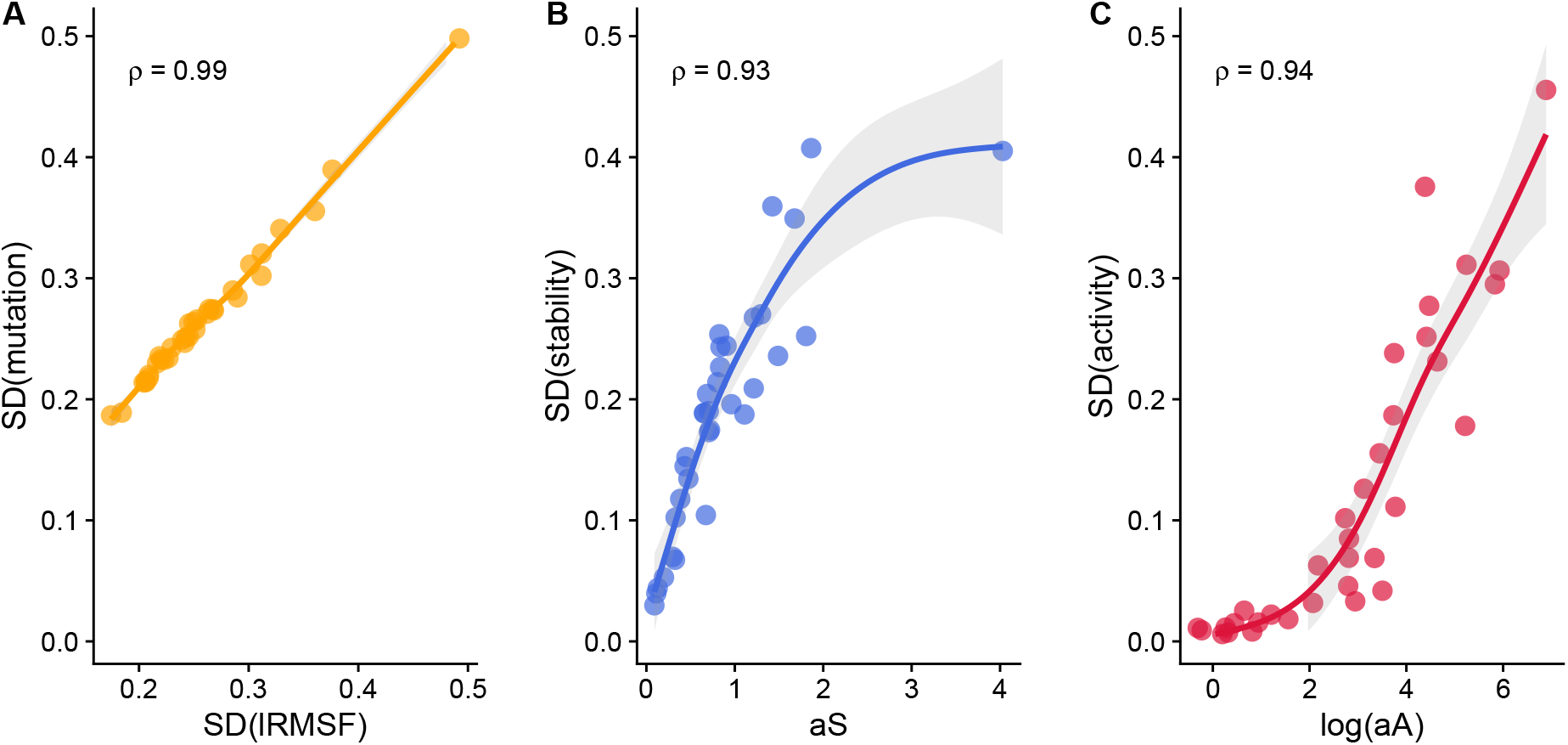
Relationship between mutation, stability, and activity contributions and underlying enzyme properties. (A) Contribution of the mutation component, SD(*ϕ*_M_), versus heterogeneity of residue flexibility, measured by the standard deviation of the logarithm of the root mean square fluctuation, SD(lRMSF). (B) Contribution of the stability component, SD(*ϕ*_S_), versus Bayesian estimate (posterior mean) of the stability selection parameter a_S_. (C) Contribution of the activity component, SD(*ϕ*_A_), versus Bayesian estimate (posterior mean) of the activity selection parameter log(a_A_). Each point represents one enzyme family. Loess fits (lines) with confidence intervals (gray shading) and Spearman correlation coefficients are shown.

The strong correlation between the mutation contribution and flexibility heterogeneity is expected because mutational structural effects correlate closely with residue flexibility (Marcos and Echave, 2020), so the mutation contribution tracks flexibility heterogeneity, an architectural property of the protein. For the stability and activity contributions, the strong correlations with a_S_ and a_A_ show that most of their variation across enzyme families reflects differences in selection strengths rather than differences in the distributions of ΔΔ*G* and ΔΔ*G*^‡^ across residues.

## 3 Discussion

We extended the MSA model (Echave, 2019) to predict residue-dependent structural divergence profiles in enzyme evolution. In MSA, mutations perturb the protein structure and fix with probabilities that depend on their effects on stability (ΔΔG, controlled by a_S_) and catalytic activity (ΔΔG^*‡*^, controlled by a_A_). The predicted divergence profile is a weighted average of mutational displacements, where the weights are fixation probabilities. Without selection, the profile is determined entirely by the protein’s mechanical response to mutation, which varies across residues; turning on a_S_ and a_A_ reweights the ensemble, modifying the profile. Turning each selection parameter on or off allows us to test whether each constraint is needed to account for the observations, and to isolate how much each one contributes to the predicted profile.

We applied MSA to 34 enzyme families and found that the model recapitulates observed structural divergence profiles, and that mutation, stability, and activity constraints each contribute. On average, mutation constraints account for roughly half the profile, with the remainder divided between stability and activity constraints. This division, however, varies widely across families: while mutation is consistently the largest contributor, stability and activity can each range from a minor to the dominant contribution.

We next asked what determines the variation of these components among enzyme families. Given the model’s structure, variation can only come from differences in the distributions of mutational effects, which are internal properties of each protein’s architecture, or in the selection parameters a_S_ and a_A_, which reflect external selective pressures, or both. The mutation contribution tracks flexibility heterogeneity, which is purely an architectural property. The stability and activity contributions depend both on their respective selection parameters and on the distributions of ΔΔG and ΔΔG^*‡*^ across residues. In practice, however, most of the variation among enzyme families in the stability and activity contributions is accounted for by variation in a_S_ and a_A_ alone. Thus, protein architecture determines the mutational effects that generate all three components, but family-to-family differences in the stability and activity contributions are mainly set by how strongly each enzyme family is selected for stability and activity.

How well does the model perform? MSA explains on average 44% of the variance in observed structural divergence profiles. For context, the best structure-based predictors of the analogous problem for sequence evolution (variation of substitution rates among residues) explain 30–42% of the variance (Jack et al., 2016; Nagar et al., 2022), despite decades of work on that problem (see (Echave et al., 2016) for a review). Part of the remaining variance likely reflects observational noise from sources such as limited numbers of homologues, alignment errors, or variability in the experimental conditions under which structures were determined. Part may reflect constraints not included in the model; although we studied single-domain monomeric enzymes to minimise complications from protein–protein interactions, cofactor binding, or allosteric effects, these cannot be entirely excluded. The model could be improved by incorporating additional constraints, but even in its current form, it can serve as a null model, both as a benchmark for further model development and as a tool to identify missing constraints. Families where MSA systematically underperforms, or residues that are clear outliers within otherwise well-fitted profiles, would point to constraints not considered by the current model.

Beyond improving the model or using it as a null model to find additional constraints, the present work raises two fundamental questions about the variation of a_S_ and a_A_ among enzyme families: what are its consequences, and what are its causes.

The first question is what further patterns the variation of a_S_ and a_A_ among enzyme families may explain. In the present work, we used the MSA model to study the variation of structural divergence among residues within families. A natural next step is to ask whether it also accounts for differences in the rate of structural divergence among families. This is a long-standing open problem: the rate of structural divergence varies several-fold among families, but no explanation has been found in protein structural class, functional category, or rate of sequence evolution (Wood and Pearson, 1999; Illergård et al., 2009). The MSA model predicts not only residue-level rates but also the protein-level rate of structural divergence, and both depend on a_S_ and a_A_. Because these parameters vary widely among families, MSA predicts that they should produce differences in rates of structural divergence, potentially solving this open problem. This would also link two problems that have been treated as separate: why divergence varies among residues within a protein, and why it varies among proteins.

The second question is what determines a_S_ and a_A_. MSA provides family-specific estimates of these parameters, making it possible to look for biological correlates. For a_S_, a natural candidate is expression level. Highly expressed proteins evolve slowly (Pál et al., 2001; Drummond et al., 2006; Zhang and Yang, 2015), and the misfolding avoidance hypothesis (Drummond et al., 2005; Drummond and Wilke, 2008) attributes this to stronger selection for stability in abundant proteins, because the fitness cost of misfolded proteins scales with abundance. a_S_ should therefore correlate with expression level. For a_A_, two sources of variation can be anticipated. First, the functional optimization hypothesis (Cherry, 2010; Gout et al., 2010; Usmanova et al., 2024) proposes that highly expressed proteins also face stronger selection for function, because optimizing function allows cells to reduce the cost of expression; for enzymes, this would translate into stronger selection on catalytic efficiency, predicting that a_A_ should also correlate with expression level. Second, functional selection pressure on enzymes depends on their metabolic context. Genes encoding enzymes that catalyse high-flux reactions or reactions that cannot be bypassed through alternative pathways are under stronger purifying selection, even after controlling for enzyme abundance (Colombo et al., 2014; Aguilar-Rodríguez and Wagner, 2018). This suggests that a_A_ should correlate with metabolic flux and the availability of alternative pathways. Studying the correlations of a_S_ and a_A_ with expression level and with these metabolic properties would connect the biophysical parameters estimated here to enzyme biology. By providing family-specific estimates of these selection strengths, MSA makes it possible to ask how the molecular constraints that shape protein structure connect to the biological context in which enzymes function.

## 4 Methods

Our approach combines evolutionary modelling with biophysical calculations to predict residue-dependent structural divergence patterns. We first describe the Mutation-Stability-Activity (MSA) evolutionary framework and its integration with the Linearly Forced Elastic Network Model (LFENM) for calculating mutational effects on structure, stability, and activity. We then detail our implementation for generating structural divergence profiles, including mutational scanning, Bayesian parameter estimation, and profile prediction. Finally, we describe how we obtain observed structural divergence profiles from enzyme families and our analytical framework for nested model comparisons and constraint decomposition.

### 4.1 Theoretical framework: Mutation-Stability-Activity model

We apply the Mutation-Stability-Activity (MSA) evolutionary model Echave (2019) in combination with the Linearly Forced Elastic Network Model (LFENM) Echave (2008); Echave and Fernández (2010) to predict structural divergence patterns in enzyme evolution. This MSA-LFENM combination has been used previously to study sequence evolution Echave (2019, 2021); here we extend its application to predict structural divergence patterns.

#### Mutation-selection process

The MSA model represents evolution as a mutation-selection process consisting of repeated mutation-selection steps (McCandlish and Stoltzfus, 2014). At each evolutionary time-step, we denote the wild-type protein structure as **r**_wt_. When a mutation is introduced, it creates a mutant structure **r**_mut_ that either disappears from the population or becomes fixed, replacing the wild type. As substitutions (fixed mutations) accumulate over time, the protein structure progressively diverges from its ancestral state.

The fixation process is governed by the fixation probability function. In the MSA model, this fixation probability depends on mutational effects on protein stability and catalytic activity Echave (2019). Specifically, it is given by:

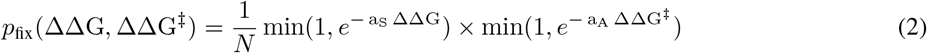

where *N* is the population size, ΔΔG = ΔG_mut_ − ΔG_wt_ is the mutational change in protein stability (negative values indicate stabilization), and ΔΔG^‡^ = ΔG^‡^_mut_ − ΔG^‡^_wt_ is the mutational change in activation energy (negative values indicate lower activation energy and, therefore, increased catalytic activity).

The parameters a_S_ and a_A_ control selection strength against destabilizing and deactivating mutations, respectively. The maximum fixation probability is 1*/N*, corresponding to neutral drift. (When mutant and wild type have equal fitness, each individual lineage has equal probability of eventual fixation, so a single mutant in a population of N has probability 1*/N* of fixing.) This maximum is achieved when mutations don’t compromise stability or activity—that is, when ΔΔG ≤ 0 and ΔΔG^‡^ ≤ 0—or when selection is absent (a_S_ = a_A_ = 0). For deleterious mutations (ΔΔG *>* 0 or ΔΔG^‡^ *>* 0), the fixation probability decreases exponentially with both the magnitude of the deleterious effect and the corresponding selection parameter.

#### Linearly Forced Elastic Network Model

To complete the evolutionary model, we need a way to calculate mutant structures and the mutational effects on stability and activation free energy. For this purpose we use the Linearly Forced Elastic Network Model (LFENM) Echave (2008).

The LFENM model represents the wild-type protein as an elastic network and mutants as perturbed elastic networks Echave (2008). The wild type protein is a network of nodes, that represent residues, connected by harmonic springs, that represent interactions. Mathematically, the wild-type’s energy is

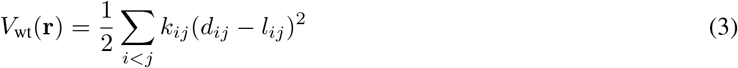

where *d*_*ij*_ is the distance between residues *i* and *j* in conformation **r**, *l*_*ij*_ is the equilibrium spring length, and *k*_*ij*_ is the spring constant. The model assumes the protein fluctuates around a single equilibrium conformation 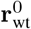. Expanding *V*_wt_ to second order around this equilibrium gives

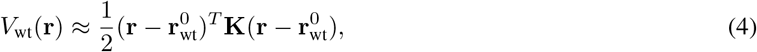

where **K** is the Hessian matrix of second derivatives of *V*_wt_ evaluated at 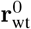.

Introducing a random mutation at a given residue is modelled by adding random perturbations to the lengths of the springs that connect the mutated site with its neighbours Echave (2008). As a result of this perturbation, there is a change of the energy landscape. The mutant’s energy is

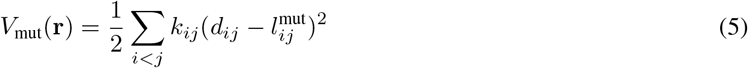

where 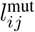 are the equilibrium spring lengths for the mutant, defined as 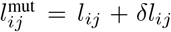 for springs involving the mutated residue, and 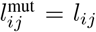 for all other springs. The perturbations *δl*_*ij*_ are independently picked from a normal distribution of mean zero and standard deviation *σ*_mut_, which is chosen small enough to ensure the perturbative approximation remains valid. (Note that this approach does not model mutations between actual amino-acid residues, but does capture the correct distribution of effects caused by actual mutations without requiring detailed side-chain modelling (Echave, 2008; Echave and Fernández, 2010; Marcos and Echave, 2020).) Because the mutation perturbs spring lengths but not force constants, expanding *V*_mut_ to second order around 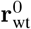 yields the same Hessian **K** plus a linear forcing term 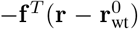, where **f** encodes the spring perturbations; this is the origin of the name “linearly forced” (see (Echave, 2008) for detailed derivations).

#### LFENM calculations of mutational effects

The LFENM model provides formulae to calculate the mutational changes of structure, stability, and activation energy. These calculations require the Hessian **K** and the force vector **f** . The force vector has the following structure: for each perturbed spring connecting the mutated residue *j* to a neighbour *i*, residue *i* experiences a force of magnitude *f*_*ij*_ = *k*_*ij*_*δl*_*ij*_ directed along the contact, and residue *j* experiences an equal and opposite reaction force. The total force on the mutated residue *j* is the vector sum of reaction forces from all its perturbed neighbours. The vector **f** is assembled from these individual force components (Echave and Fernández, 2010; Marcos and Echave, 2020). Using **K** and **f**, we now present the three key formulae.

First, we consider the calculation of the structural change, Δ**r**^0^. Minimizing *V*_mut_ with respect to **r** gives (Echave, 2008; Echave and Fernández, 2010):

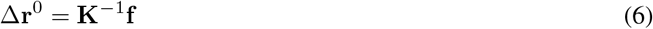

where 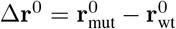 is the structural change upon mutation (**K**^−1^ denotes the pseudo-inverse, since **K** is singular due to translational and rotational invariance).

Second, we consider the calculation of the stability change, ΔΔG. Because the mutation shifts the equilibrium conformation and minimum energy but does not change the curvature of the energy well, the entropic contributions to the free energy cancel exactly, and ΔΔG reduces to the difference between the minimum energies of the mutant and wild-type potentials (see also Supplementary Section S1). This can be written Echave (2019):

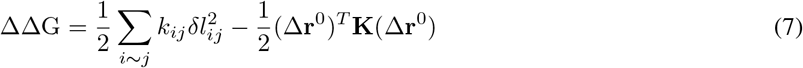

where the summation is over contacts involving the mutated site (the only non-zero *δl*_*ij*_ terms). The first term represents the local stress energy that results from perturbed contacts, while the second term accounts for the decrease in energy due to a global relaxation of the mutant’s structure.

Finally, we consider the calculation of the activation free energy change ΔΔG^‡^. We derive it here briefly; the interested reader is referred to (Echave, 2019) for full details. In transition state theory (Schowen, 1978), the reaction rate depends on the probability of the enzyme reaching the transition-state conformation, which is determined by the thermal fluctuations within the reactant-state energy well. A mutation that shifts the active-site geometry away from the transition-state conformation reduces this probability, increasing ΔG^‡^. Thus, calculating ΔΔG^‡^ requires only the reactant-state well, which is what the ENM model describes.

Following transition state theory, ΔG^‡^ can be decomposed into enzyme distortion, substrate distortion, and vertical binding energy contributions. Assuming that mutation affects only the enzyme distortion term, the other two cancel in ΔΔG^‡^. Further assuming that the wild-type active site is preorganised, meaning its native conformation is identical to the transition-state conformation, the wild-type enzyme distortion cost is zero, and ΔΔG^‡^ reduces to the cost of distorting the mutant from its equilibrium conformation to the transition-state active-site geometry. This cost is obtained from the marginal distribution of active-site coordinates, derived by integrating the mutant’s Boltzmann distribution over the non-active-site coordinates. Because the full distribution is Gaussian with covariance matrix **C** ∝ **K**^−1^, the marginal is also Gaussian, with covariance matrix **C**_*aa*_, the block of **C** corresponding to active-site coordinates. The effective stiffness of the active site, 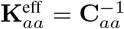, differs from the bare active-site block **K**_*aa*_ because it accounts for indirect couplings between active-site residues mediated through the rest of the protein. Using LFENM to evaluate this cost yields:

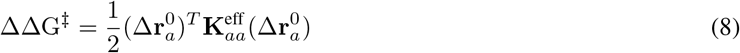

where 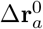 is composed by the active-site components of the structural change Δ**r**^0^.

### 4.2 Computational implementation: MSA structural divergence profiles

In this section we describe our computational implementation of MSA for predicting residue-dependent structural divergence profiles. The calculation pipeline takes a protein structure as input and proceeds through four main stages:

1. Set up the elastic network model for the wild-type structure to enable LFENM calculations of mutational effects
2. Perform a mutational scan across all residue positions, calculating how each mutation affects protein structure (Δ**r**^0^), stability (ΔΔG), and activation energy (ΔΔG^‡^) using LFENM
3. Estimate the posterior distribution of MSA model parameters (*a*_*S*_ and *a*_*A*_) from observed structural divergence data using Bayesian inference
4. Calculate MSA predicted profiles by averaging predictions across the posterior sample of parameters

#### Wild-type elastic network model setup

Specifying the elastic network model for the wild-type, amounts to specifying the parameters of the energy function shown in Eq. (3), and calculating the network matrix **K**, necessary for the calculations of mutational changes structure (Eq. (6)), stability (Eq. (7)), and activity (Eq. (8)).

There are several popular elastic network models of proteins (Hinsen, 1998; Atilgan et al., 2001; Yang et al., 2009; Ming and Wall, 2005). However, this choice is not a major issue, because LFENM predictions are very robust with respect to the specific model used (Marcos and Echave, 2020). Here, we use the parameters derived by Ming and Wall (Ming and Wall, 2005), *which are somewhat more realistic than other simpler models, and that work well for MSA predictions of sequence patterns (Echave, 2019, 2021)*. *Nodes are placed at the residue alpha carbons, C*_*α*_; spring lengths are the distances in the equilibrium native structure: 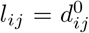, and spring force constants depend on whether contacts are along the sequence or not: *k*_*ij*_ = 189 kcal/mol for sequence neighbours, and *k*_*ij*_ = 4.5 kcal/mol for other contacts within cut-off distance *R*_0_ = 10.5 Å .

#### Mutational scanning calculations

After setting up the wild-type elastic network model, we perform a mutational scan. For each residue *j* we generate ten independent mutants (*m* = 1, …, 10). Each mutant is produced by perturbing the lengths of the springs connected to residue *j*, excluding its sequence neighbours (*j* ± 1) that represent the covalent backbone. The perturbations *δl*_*ij*_ are drawn independently from a normal distribution with mean 0 and standard deviation *σ*_mut_ = 0.3 Å . This amplitude is small enough to avoid unphysical conformations, and its exact value does not affect results because our structural-divergence measures remove any uniform scaling.

For every mutation (*j, m*) we compute the corresponding force vector **f** and then evaluate the mutational effects: the structural change Δ**r**^0^(*j, m*) (Eq. (6)), the stability change ΔΔG(*j, m*) (Eq. (7)), and the activation-energy change ΔΔG^‡^(*j, m*) (Eq. (8)).

This procedure samples the typical range of structural, stability, and activation-energy changes caused by amino-acid substitutions without requiring explicit sequence information or detailed side-chain modelling.

#### Bayesian parameter estimation

We infer the stability and activity selection strengths, a_S_ and a_A_, using a Bayesian framework.

#### Forward model

As a first step we specify a forward model, a computational procedure that predicts the residue-dependent structural-divergence profile for any given pair of parameter values.

To efficiently compute structural divergence profiles for different parameter values, we exploit a key property of evolutionary structural divergence: as substitutions accumulate over evolutionary time, the number of substitutions scales all residue RMSD values uniformly by the same factor *F*(*n*), where *n* is the total number of substitutions. This uniform scaling makes the ratios RMSD(*i*)*/* RMSD(*j*) independent of *n*, since the scaling factor cancels out. Since these ratios fully determine the relative pattern of structural divergence among residues, which is all we seek to predict, we need only consider *n* = 1. Instead of simulating multi-substitution evolutionary trajectories for every trial pair of parameter values, we perform the mutation scan once, independently of a_S_ and a_A_, to obtain the mutation-induced displacement vectors Δ**r**^0^(*j, m*), and for each parameter pair simply compute weighted averages of these pre-computed displacements as described below.

For a given pair of parameter values (a_S_, a_A_) we calculate the fixation probability *p*_fix_(*j, m*; a_S_, a_A_) of every mutant (*j, m*) using Eq. (2), together with the mutational effects ΔΔG(*j, m*) and ΔΔG^‡^(*j, m*) obtained in the mutation-scan step. These probabilities are then normalised to give the relative abundance of each mutant structure in the evolved ensemble:

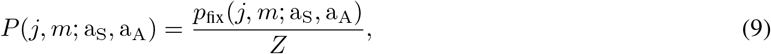

where the normalising constant *Z* is the sum of *p*_fix_ over all mutants.

From this ensemble composition we calculate the expected structural divergence. For each residue *i*, the root-meansquare deviation is the probability-weighted displacement:

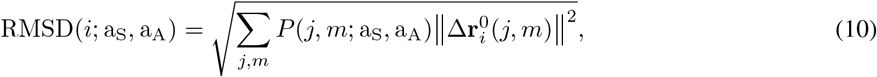

where 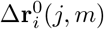 is the three-dimensional mutation-induced displacement vector of residue *i* obtained in the mutational-scan step.

To focus on residue-to-residue variation we use the centred log–root-mean-square displacement:

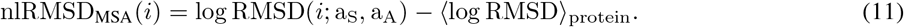

Equations (9)–(11) together constitute the forward model, mapping any pair of parameters (a_S_, a_A_) to the predicted profile of centred log–RMSD values.

#### Bayesian inference

To estimate the parameters we combine the forward model with the data using Bayes’ rule, giving the posterior distribution of parameters:

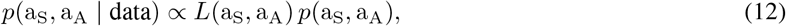

where *L*(a_S_, a_A_) is the likelihood of the observed centred log-RMSD profile. Assuming normally distributed residuals, the log-likelihood is given by:

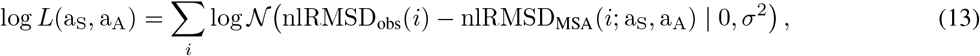

where 𝒩 (· | 0, *σ*^2^) is the density of a normal distribution with mean zero and variance *σ*^2^.

We assigned weakly informative uniform priors:

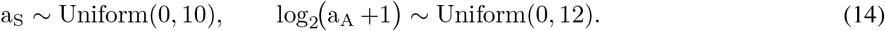

Stability selection strengths across enzyme families vary within roughly one order of magnitude, whereas activity selection spans several orders Echave (2019, 2021). Accordingly we place a direct uniform prior on a_S_, while for a_A_ we work on a logarithmic scale. The transformation log_2_(a_A_ +1) keeps the mapping defined even when a_A_ is near zero.

Posterior sampling uses a Metropolis-Hastings Markov chain Monte Carlo algorithm. At each step the algorithm proposes a new pair of parameters (a_S_, a_A_) and accepts or rejects it based on the ratio of posterior probabilities of the proposed and current states. The prior is uniform on a_S_ ∈ [0, 10] and log_2_(a_A_ +1) ∈ [0, 12], so proposals outside these ranges are rejected automatically and within them acceptance is determined by the likelihood ratio alone. The resulting Markov chain provides the posterior sample of (a_S_, a_A_) used for subsequent analyses. We ran 2,500 iterations with a 500-iteration burn-in and checked convergence in runs extended to 10,000 iterations.

Throughout Results and figures, reported values of a_S_ and a_A_ correspond to posterior means from the MCMC sample, unless otherwise noted.

#### Posterior-averaged predictions

Having obtained an MCMC sample from the posterior of (a_S_, a_A_), we calculate the model’s residue-dependent structural-divergence profile by averaging predictions over these sampled parameter values:

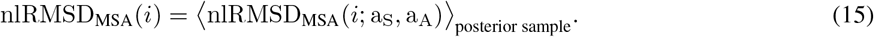

This posterior-sample mean constitutes the MSA model’s primary output for all subsequent analyses.

### 4.3 Observed structural divergence profiles

This section describes how we obtain observed residue-dependent structural divergence profiles from evolutionary data, to serve as targets for MSA model validation. We describe the enzyme families dataset, structural alignment and superposition procedures, and quantification of residue-level structural divergence.

#### Dataset of enzyme families

We used a dataset of 34 families of functionally conserved, monomeric, single-domain enzymes, previously curated from the Mechanism and Catalytic Site Atlas (M-CSA) database (Echave and Carpentier, 2025). Each family consists of a reference enzyme (for which mechanism and active site information has been experimentally determined) and a set of homologues. The dataset construction process, detailed in our previous work, applied sequential filtering to ensure functional conservation and structural quality, focusing on monomeric, single-domain proteins to avoid complications from inter-domain or inter-chain interactions. The final dataset comprises 34 enzyme families with broad structural and functional diversity: 27 *α/β*, 5 all-*β*, and 2 all-*α* folds across major CATH architectures. Functionally, hydrolases dominate (20 families), with transferases, lyases, isomerases, and oxidoreductases also represented. Family sizes range from 4–23 proteins, with reference enzymes spanning 98–447 residues (Supplementary Tables S1 and S2).

#### Alignment and superposition

Homologous sequences were aligned using MUSCLE v5 Edgar (2022) via the bio3d R package Grant et al. (2021). We chose sequence over structural alignment because functionally conserved families retain sufficient sequence similarity for reliable alignment while providing an alignment basis independent of structural considerations for studying structural divergence. Following sequence alignment, protein structures were superimposed by minimizing the global (protein-level) RMSD of C*α* coordinates, with each homologue aligned to its family’s reference enzyme.

#### Calculation of observed structural divergence profiles

Structural divergence was quantified at each residue position using the residue-level root mean square deviation of C*α* coordinates across homologous structures. For residue *i* in a family of *N*_hom_ homologous structures:

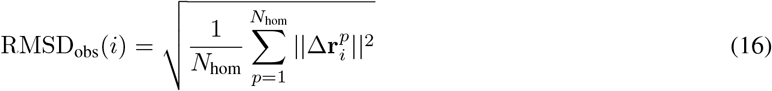

where 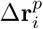 is the deviation of residue *i* in structure *p* from its average position.

We applied logarithmic transformation and centring to match the format used for model predictions:

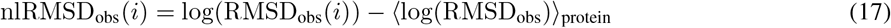

where the second term is the mean log-RMSD across all residues in the protein. These centred log-RMSD profiles serve as the observational targets for model validation and parameter fitting.

### 4.4 Analysis

Once we have obtained observed and model structural divergence profiles, we analysed them in the following four steps:

1. Model validation by comparison with observed structural divergence profiles
2. Nested model comparison to assess whether each constraint contributes to explaining the data
3. Decomposition analysis to quantify the relative importance of different constraints
4. Identification of enzyme properties that determine why constraint importance varies among families

#### Model validation

We validate MSA predictions by comparing them with observed structural divergence profiles and with predictions from our previous empirical model M12 (Echave and Carpentier, 2025). M12 is a shape-constrained generalized additive model that smooths observed structural divergence by fitting it to a fucnction that is the sum of two components: a function of residue fexibility plus a function of distance to the active site. M12 provides a flexible fit that effectively removes observational noise, serving as a benchmark for the underlying signal. Model performance is quantified using Pearson correlation coefficients between predicted and observed profiles across all enzyme families. We also calculate correlations between MSA and M12 predictions to assess whether the parsimonious mechanistic approach captures similar patterns to the flexible empirical model.

#### Nested model comparison

To determine whether all three MSA constraints influence structural divergence patterns, we examine four nested model variants that systematically add components: M0, MM, MS, and MSA. The baseline null model M0 represents uniform structural divergence across all residues, which corresponds to nlRMSD_M0_(*i*) = 0 since we use centred profiles. The mutation model MM includes only mutational effects by setting both selection parameters to zero. The mutation-stability model MS adds stability selection by setting only the activity selection parameter to zero. The full MSA model includes all three constraints.

To obtain predictions from each nested model, we use the posterior sample of parameters from the full MSA fit. For each parameter pair in this sample, we calculate the corresponding nested model prediction by setting the appropriate parameters to zero (e.g., a_S_ = a_A_ = 0 for MM, or a_A_ = 0 for MS). We then average these predictions over the entire posterior sample. This approach avoids potential compensation effects that could arise from refitting simplified models where remaining parameters might inflate to compensate for missing constraints.

We quantify the explanatory power of each model using explained deviance *D*^2^:

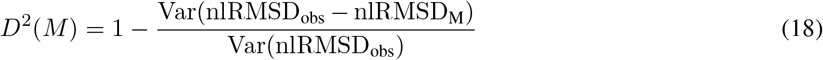

where *M* represents the model being evaluated, *D*^2^ = 0 for the baseline model and *D*^2^ = 1 for perfect fit. We test the contribution of each constraint by examining the increments Δ*D*^2^ in the progression M0 → MM → MS → MSA. Positive increments at each step would indicate that the added constraint contributes explanatory power beyond what the simpler models already capture.

#### Decomposition analysis

To isolate the contribution of each constraint, we decompose the MSA prediction at each residue position into three components:

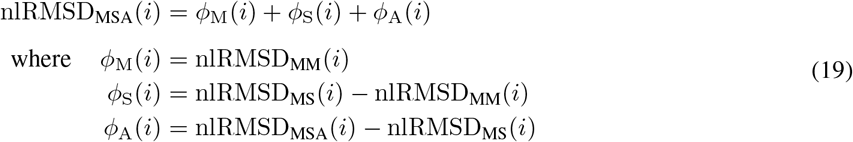

Here *i* denotes the residue position, and the components represent the mutation-only contribution, the additional effect of stability selection, and the additional effect of activity selection, respectively.

This decomposition approach is necessary because Δ*D*^2^ values from the nested comparison, while establishing that constraints contribute, cannot isolate the individual contribution of each constraint due to interaction effects.

Two problems arise with variance-based measures like *D*^2^. First, *D*^2^ variation among enzymes depends on observational noise, not just model accuracy. Second, interaction effects contaminate the measures: since *D*^2^ is based on squared deviations, interaction terms arise between components. (For example, from the definitions above nlRMSD_MS_(*i*) = *ϕ*_M_(*i*) + *ϕ*_S_(*i*), so *D*^2^(MS) = *D*^2^(*ϕ*_M_) + *D*^2^(*ϕ*_S_) + 2Cor(*ϕ*_M_, *ϕ*_S_), where Cor denotes correlation. The increment Δ*D*^2^ = *D*^2^(MS) − *D*^2^(MM) therefore includes an interaction term, preventing Δ*D*^2^ from measuring the stability constraint’s contribution cleanly.)

To avoid these issues, we quantify component contributions using the standard deviations of the individual components across residue positions, SD(*ϕ*_M_), SD(*ϕ*_S_), and SD(*ϕ*_A_), which directly measure each constraint’s contribution to profile variation without interaction effects. Relative contributions are calculated by dividing each component’s standard deviation by the sum of all component standard deviations.

#### Determinants of constraint contributions

We examine which enzyme properties explain the variation in SD(*ϕ*_M_), SD(*ϕ*_S_), and SD(*ϕ*_A_) across enzyme families.

For mutation contributions, previous work shows that mutational structural effects correlate strongly with protein flexibility patterns Marcos and Echave (2020). We therefore examine the correlation between SD(*ϕ*_M_) and flexibility heterogeneity SD(lRMSF), where lRMSF(*i*) is the logarithm of the root mean square fluctuation of residue *i*, which is obtained using the reference protein’s ENM.

For stability and activity contributions, previous findings indicate that selection contributions depend on their respective selection strengths Echave (2021). We therefore examine correlations between SD(*ϕ*_S_) and the stability selection parameter a_S_, and between SD(*ϕ*_A_) and the activity selection parameter a_A_.

We use Spearman correlation coefficients to assess these relationships.

## 5 Data availability

All processed data and code needed to reproduce the figures and tables in this paper are available at https://github.com/jechave/msa_profiles_shared.

## Acknowledgments

This work was supported by CONICET (grant PIP-11220210100462) and regular ISYEB funding.

