## Supplementary document for "Why structural divergence varies among residues in enzyme evolution: contributions of mutation, stability, and activity constraints"

#### S1 Toy model of mutational effects on structure, stability, and activation energy

This section derives the mutational effects on structure, stability, and activation energy for a minimal model: a single active-site residue connected to two regions of the protein scaffold (Figure~S1). We then show that the LFENM matrix formalism (Eqs.~4–6 of the main text) reproduces the results.

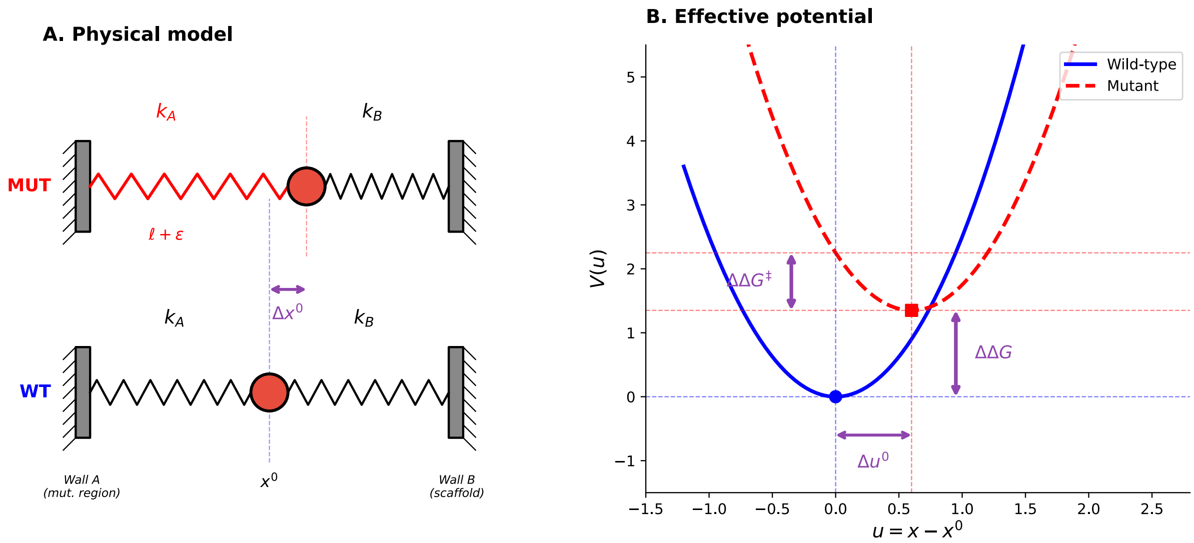


Figure S1: **Toy illustration of how LFENM calculates mutational effects on structure, stability, and activation energy.** (A) Physical model. A single residue (red circle) is connected by harmonic springs to the mutation region (Wall A, spring constant $k_A$) and the scaffold (Wall B, spring constant $k_B$). Bottom: wild-type equilibrium at $x^0$. Top: a mutation perturbs the natural length of the $k_A$ spring by $\varepsilon$ (red), shifting the equilibrium by $\Delta x^0$. (B) Effective potential energy as a function of displacement $u = x - x^0$. The wild-type potential (blue, solid) is minimised at $u = 0$; the mutant potential (red, dashed) is minimised at $u = \Delta u^0$. The vertical arrows indicate $\Delta\Delta G$ (stability cost) and $\Delta\Delta G^{\ddagger}$ (activation energy cost).

##### Setup

Consider a single bead (representing an active-site residue) at position $x$, connected by two harmonic springs to fixed walls representing rigid parts of the protein scaffold (Figure~S1A). Wall~A, at position $A$, represents the region of the protein containing the mutation site; Wall~B, at position $B$, represents the rest of the scaffold. The springs have force constants $k_A$ and $k_B$ and natural lengths $\ell_A = x^0 - A$ and $\ell_B = B - x^0$, where $x^0$ is the wild-type equilibrium position.

The wild-type potential energy is

$$V_{\mathrm{wt}} = \frac{k_A}{2}(x - A - \ell_A)^2 + \frac{k_B}{2}(B - x - \ell_B)^2. \tag{S1}$$

Defining the displacement from the wild-type equilibrium, $u = x - x^0$, we obtain:

$$V_{\text{wt}} = \frac{k_A + k_B}{2}\, u^2. \tag{S2}$$

The wild-type minimum is at $u = 0$ with $V_{\text{wt}}(0) = 0$.

#### Mutation

A mutation in region~A is modelled as a perturbation $\varepsilon$ to the natural length of the $k_A$ spring: $\ell_A \to \ell_A + \varepsilon$. The mutant potential is

$$V_{\text{mut}} = \frac{k_A}{2}(u - \varepsilon)^2 + \frac{k_B}{2}\, u^2, \tag{S3}$$

which can be rewritten as

$$V_{\text{mut}} = \frac{k_A + k_B}{2}\, u^2 - k_A\varepsilon\, u + \frac{k_A}{2}\,\varepsilon^2. \tag{S4}$$

The mutation adds a linear force $k_A\varepsilon$ pulling the bead toward a new position, plus a constant stress energy $k_A\varepsilon^2/2$. The quadratic term $(k_A + k_B)/2$ is unchanged: the mutation shifts the parabola horizontally and vertically but does not change its curvature (Figure~S1B). This is because the mutation perturbs the spring length, not the spring constant, so the Hessian is identical for wild type and mutant.

#### Structural change

Minimising $V_{\text{mut}}$ with respect to $u$:

$$(k_A + k_B)\, u = k_A\, \varepsilon, \tag{S5}$$

the mutant equilibrium displacement is

$$\Delta u^0 = \frac{k_A\, \varepsilon}{k_A + k_B}. \tag{S6}$$

The bead shifts toward the position favoured by the mutated spring, but the opposing spring $k_B$ prevents it from reaching $u = \varepsilon$. The compromise position $\Delta u^0$ lies between 0 and $\varepsilon$: closer to 0 when the coupling to the scaffold is strong ($k_B \gg k_A$) and closer to $\varepsilon$ when the coupling to the mutation region is strong ($k_A \gg k_B$).

#### Stability change ($\Delta\Delta G$)

Because the Hessian is the same for wild type and mutant, the shape of the potential is unchanged by the mutation. The thermal fluctuations around each minimum are therefore identical, and the entropic contributions to the free energy cancel exactly. The free energy difference between mutant and wild type thus reduces to the difference in potential energy at their respective minima:

$$\Delta\Delta G = V_{\text{mut}}(\Delta u^0) - V_{\text{wt}}(0). \tag{S7}$$

Evaluating $V_{\text{mut}}$ at $\Delta u^0$:

$$\Delta\Delta G = \frac{k_A}{2}(\Delta u^0 - \varepsilon)^2 + \frac{k_B}{2}(\Delta u^0)^2. \tag{S8}$$

Substituting Eq.~(S6):

$$\Delta\Delta G = \frac{k_A\, k_B}{2(k_A + k_B)}\,\varepsilon^2. \tag{S9}$$

This can be decomposed as

$$\Delta\Delta G = \underbrace{\frac{k_A}{2}\,\varepsilon^2}_{\text{stress}} - \underbrace{\frac{k_A^2}{2(k_A + k_B)}\,\varepsilon^2}_{\text{relaxation}}. \tag{S10}$$

The stress term $k_A\varepsilon^2/2$ is the energy cost if the bead were frozen at its wild-type position ($u = 0$). The relaxation term is the energy recovered by allowing the bead to shift to its new equilibrium. The stability cost $\Delta\Delta G$ is the residual stress that cannot be relaxed away: it arises from the conflict between the mutated and unmutated springs, which cannot be simultaneously satisfied.

#### Activation energy change ($\Delta\Delta G^{\ddagger}$)

The wild-type active-site geometry ($u = 0$) is assumed to be optimal for catalysis, meaning it coincides with the transition-state conformation. In the mutant, the active site has relaxed to $u = \Delta u^0$. To adopt the catalytically competent conformation, the mutant must be distorted from $\Delta u^0$ to $u = 0$. The cost of this conformational distortion is (Figure~S1B):

$$\Delta\Delta G^{\ddagger} = V_{\text{mut}}(0) - V_{\text{mut}}(\Delta u^0). \tag{S11}$$

Substituting Eq.~(S3) and Eq.~(S6):

$$\Delta\Delta G^{\ddagger} = \frac{k_A}{2}\,\varepsilon^2 - \frac{k_A\,k_B}{2(k_A+k_B)}\,\varepsilon^2 = \frac{k_A^2}{2(k_A+k_B)}\,\varepsilon^2. \tag{S12}$$

#### Connection to the LFENM matrix formalism

The derivations above used direct minimisation of the potential. We now show that the LFENM matrix formalism reproduces all three results. In the full LFENM, a mutation at residue $j$ is encoded as a force vector $\mathbf{f}$, and the three mutational effects are calculated from the network matrix $\mathbf{K}$ (the Hessian of the wild-type energy) and its pseudo-inverse, the variance-covariance matrix $\mathbf{C}$:

$$\Delta\mathbf{r}^0 = \mathbf{C}\mathbf{f}, \tag{S13}$$

$$\Delta\Delta G = \frac{1}{2}\sum_{i\sim j} k_{ij}\,\delta l_{ij}^2 - \frac{1}{2}(\Delta\mathbf{r}^0)^T\mathbf{K}\,\Delta\mathbf{r}^0, \tag{S14}$$

$$\Delta\Delta G^{\ddagger} = \frac{1}{2}(\Delta\mathbf{r}_a^0)^T\mathbf{K}_{aa}^{\text{eff}}\,\Delta\mathbf{r}_a^0, \tag{S15}$$

where $\Delta\mathbf{r}_a^0$ is the active-site subset of $\Delta\mathbf{r}^0$ and $\mathbf{K}_{aa}^{\text{eff}} = \mathbf{C}_{aa}^{-1}$ is the inverse of the active-site block of $\mathbf{C}$.

For the one-bead model, $\mathbf{K}$, $\mathbf{C}$, and $\mathbf{f}$ reduce to scalars:

$$K = \frac{d^2V_{\text{wt}}}{du^2} = k_A + k_B, \qquad C = K^{-1} = \frac{1}{k_A+k_B}, \qquad f = k_A\,\varepsilon. \tag{S16}$$

**Structural change.** Eq.~(S13) gives:

$$\Delta u^0 = C\cdot f = \frac{k_A\,\varepsilon}{k_A+k_B}, \tag{S17}$$

confirming Eq.~(S6).

**Stability change.** Eq.~(S14) gives:

$$\Delta\Delta G = \frac{1}{2}k_A\varepsilon^2 - \frac{1}{2}(\Delta u^0)^2K = \frac{k_A}{2}\,\varepsilon^2 - \frac{k_A^2\varepsilon^2}{2(k_A+k_B)}, \tag{S18}$$

confirming Eq.~(S10).

**Activation energy change.** In the toy model there is only one degree of freedom, the active-site coordinate $u$, so the active-site subspace is the entire space and no marginalisation over non-active-site coordinates is needed. Consequently $C_{aa} = C = 1/(k_A+k_B)$ and $K_{aa}^{\text{eff}} = C_{aa}^{-1} = k_A + k_B$. Eq.~(S15) with $\Delta\mathbf{r}_a^0 = \Delta u^0$ gives:

$$\Delta\Delta G^{\ddagger} = \frac{1}{2}(\Delta u^0)^2\,(k_A+k_B) = \frac{k_A^2\varepsilon^2}{2(k_A+k_B)}, \tag{S19}$$

confirming Eq.~(S12). In a multi-residue protein, marginalising over non-active-site coordinates generally yields $\mathbf{K}_{aa}^{\text{eff}} \neq \mathbf{K}_{aa}$, because $\mathbf{K}_{aa}^{\text{eff}}$ captures indirect couplings between active-site residues mediated through the rest of the protein; their equality here is a special feature of the one-bead model.

### S2 Supplementary Tables

Table S1: Dataset entries and reference protein

| M-CSA | PDB | Name | Source | Family |
|---|---|---|---|---|
| 2 | 1btl_A | Beta-lactamase TEM | Escherichia coli | Class-A beta-lactamase family |
| 15 | 1znb_A | Metallo-beta-lactamase type 2 | Bacteroides fragilis | Metallo-beta-lactamase superfamily |
| 71 | 1eug_A | Uracil-DNA glycosylase | Escherichia coli B | Uracil-DNA glycosylase (UDG) superfamily |
| 98 | 1bsz_C | Peptide deformylase | Escherichia coli K-12 | Polypeptide deformylase family |
| 109 | 1d3g_A | Dihydroorotate dehydrogenase (quinone), mitochondrial | Homo sapiens | Dihydroorotate dehydrogenase family |
| 148 | 1onr_A | Name not found | Escherichia coli | Transaldolase family |
| 151 | 1q0n_A | 2-amino-4-hydroxy-6-hydroxymethyldihydropteridine pyrophosphokinase | Escherichia coli | HPPK family |
| 160 | 1ako_A | Exodeoxyribonuclease III | Escherichia coli K-12 | DNA repair enzymes AP/ExoA family |
| 164 | 1ruv_A | Ribonuclease pancreatic | Bos taurus | Pancreatic ribonuclease family |
| 174 | 9pap_A | Papain | Carica papaya | Peptidase C1 family |
| 216 | 1ca2_A | Carbonic anhydrase 2 | Homo sapiens | Alpha-carbonic anhydrase family |
| 252 | 1igs_A | Indole-3-glycerol phosphate synthase | Saccharolobus solfataricus | TrpC family |
| 257 | 1xx2_A | Beta-lactamase | Enterobacter cloacae | Class-C beta-lactamase family |
| 258 | 1sml_A | Metallo-beta-lactamase L1 type 3 | Stenotrophomonas maltophilia | Metallo-beta-lactamase superfamily |
| 290 | 1zio_A | Adenylate kinase | Geobacillus stearothermophilus | Adenylate kinase family |
| 328 | 1lbm_A | N-(5&apos | Thermotoga maritima | TrpF family |
| 351 | 1snz_A | Galactose mutarotase | Homo sapiens | Aldose epimerase family |
| 362 | 1d6o_A | Peptidyl-prolyl cis-trans isomerase FKBP1A | Homo sapiens | FKBP-type PPIase family |
| 376 | 1a4l_A | Adenosine deaminase | Mus musculus | Metallo-dependent hydrolases superfamily |
| 394 | 1aj0_A | Dihydropteroate synthase | Escherichia coli | DHPS family |
| 444 | 1cel_A | Exoglucanase 1 | Trichoderma reesei | Glycosyl hydrolase 7 (cellulase C) family |
| 462 | 1pnt_A | Low molecular weight phosphotyrosine protein phosphatase | Bos taurus | Low molecular weight phosphotyrosine protein phosphatase family |
| 467 | 1cv2_A | Haloalkane dehalogenase | Sphingomonas paucimobilis | Haloalkane dehalogenase family |
| 480 | 1czf_A | Endopolygalacturonase II | Aspergillus niger | Glycosyl hydrolase 28 family |
| 597 | 1uch_A | Ubiquitin carboxyl-terminal hydrolase isozyme L3 | Homo sapiens | Peptidase C12 family |
| 681 | 1gq8_A | Pectinesterase | Daucus carota | Pectinesterase family |
| 693 | 2rnf_A | Ribonuclease 4 | Homo sapiens | Pancreatic ribonuclease family |
| 749 | 1nml_A | No information found | Marinobacter nauticus | No information found |
| 814 | 1glo_A | Cathepsin S | Homo sapiens | Peptidase C1 family |
| 858 | 1mrq_A | Aldo-keto reductase family 1 member C1 | Homo sapiens | Aldo/keto reductase family |
| 877 | 1pbg_A | 6-phospho-beta-galactosidase | Lactococcus lactis | Glycosyl hydrolase 1 family |
| 908 | 1rtu_A | Ribonuclease U2 | Ustilago sphaerogena | Ribonuclease U2 family |
| 923 | 2acy_A | Acylphosphatase-1 | Bos taurus | Acylphosphatase family |
| 931 | 2pth_A | Peptidyl-tRNA hydrolase | Escherichia coli K-12 | PTH family |

Table S2: Dataset properties

| M-CSA | N | <Id%> | <RMSD> | Length | AS size | AS RMSD | CATH | EC |
|---|---|---|---|---|---|---|---|---|
| 2 | 6 | 43 | 1.04 | 263 | 6 | 0.35 | alpha/beta | Hydrolases |
| 15 | 6 | 35 | 1.16 | 225 | 8 | 0.40 | alpha/beta | Hydrolases |
| 71 | 8 | 43 | 1.08 | 223 | 4 | 0.53 | alpha/beta | Hydrolases |
| 98 | 10 | 37 | 2.15 | 168 | 7 | 0.44 | alpha/beta | Hydrolases |
| 109 | 4 | 44 | 1.31 | 359 | 7 | 1.78 | alpha/beta | Oxidoreductases |
| 148 | 5 | 53 | 1.06 | 316 | 5 | 0.57 | alpha/beta | Transferases |
| 151 | 4 | 47 | 1.14 | 158 | 4 | 1.08 | alpha/beta | Transferases |
| 160 | 10 | 34 | 1.49 | 256 | 7 | 0.38 | alpha/beta | Hydrolases |
| 164 | 5 | 48 | 1.09 | 123 | 5 | 0.50 | alpha/beta | Lyases |
| 174 | 12 | 43 | 1.01 | 212 | 4 | 0.75 | alpha/beta | Hydrolases |
| 216 | 5 | 39 | 1.09 | 256 | 6 | 0.30 | alpha/beta | Lyases |
| 252 | 6 | 35 | 1.75 | 245 | 7 | 0.61 | alpha/beta | Lyases |
| 257 | 10 | 45 | 0.94 | 359 | 6 | 0.43 | alpha/beta | Hydrolases |
| 258 | 4 | 38 | 1.97 | 263 | 7 | 0.87 | alpha/beta | Hydrolases |
| 290 | 7 | 47 | 1.10 | 217 | 5 | 0.55 | alpha/beta | Transferases |
| 328 | 4 | 33 | 1.47 | 193 | 2 | 0.64 | alpha/beta | Isomerases |
| 351 | 4 | 33 | 1.36 | 341 | 3 | 0.47 | all-beta | Isomerases |
| 362 | 7 | 46 | 0.92 | 107 | 6 | 0.43 | alpha/beta | Isomerases |
| 376 | 4 | 44 | 1.30 | 349 | 6 | 0.37 | alpha/beta | Hydrolases |
| 394 | 10 | 40 | 2.02 | 280 | 2 | 0.59 | alpha/beta | Transferases |
| 444 | 6 | 46 | 1.19 | 431 | 4 | 0.44 | all-beta | Hydrolases |
| 462 | 6 | 37 | 1.15 | 155 | 6 | 0.34 | alpha/beta | Hydrolases |
| 467 | 5 | 45 | 1.03 | 293 | 5 | 0.45 | alpha/beta | Hydrolases |
| 480 | 4 | 44 | 1.05 | 335 | 6 | 0.42 | all-beta | Hydrolases |
| 597 | 4 | 38 | 1.76 | 204 | 4 | 0.90 | alpha/beta | Hydrolases |
| 681 | 4 | 31 | 2.88 | 300 | 5 | 0.80 | all-beta | Hydrolases |
| 693 | 7 | 38 | 1.62 | 120 | 3 | 0.49 | alpha/beta | Hydrolases |
| 749 | 6 | 56 | 1.96 | 316 | 1 | 5.17 | all-alpha | NA |
| 814 | 4 | 46 | 0.72 | 215 | 4 | 0.37 | alpha/beta | Hydrolases |
| 858 | 16 | 38 | 1.88 | 322 | 4 | 0.68 | alpha/beta | Oxidoreductases |
| 877 | 23 | 37 | 1.52 | 447 | 2 | 0.48 | alpha/beta | Hydrolases |
| 908 | 5 | 45 | 1.45 | 107 | 4 | 0.49 | alpha/beta | Lyases |
| 923 | 4 | 34 | 0.99 | 98 | 2 | 0.49 | alpha/beta | Hydrolases |
| 931 | 7 | 39 | 1.40 | 193 | 4 | 0.43 | alpha/beta | Hydrolases |

N: number of family members; <Id%>: average family seq. identity; <RMSD>: average RMSD over all residues; Length: number of residues of reference protein; AS size: number of active-site residues; AS RMSD: average RMSD over active-site residues; CATH: CATH class; EC: EC class

Table S3: MSA model parameters and SHAP decomposition

| M-CSA | PDB | $a_S$ | $a_A$ | $D^2$ | $R$ | sd($\phi_{\text{mut}}$) | sd($\phi_{\text{stab}}$) | sd($\phi_{\text{act}}$) |
|---|---|---|---|---|---|---|---|---|
| 2 | 1btl_A | 0.20 | 87.50 | 0.49 | 0.67 | 0.23 | 0.05 | 0.28 |
| 15 | 1znb_A | 0.47 | 42.43 | 0.65 | 0.78 | 0.34 | 0.13 | 0.24 |
| 71 | 1eug_A | 0.91 | 2.56 | 0.75 | 0.83 | 0.39 | 0.24 | 0.02 |
| 98 | 1bsz_C | 4.03 | 33.30 | 0.81 | 0.82 | 0.36 | 0.41 | 0.04 |
| 109 | 1d3g_A | 0.82 | 0.73 | 0.44 | 0.62 | 0.26 | 0.25 | 0.01 |
| 148 | 1onr_A | 0.69 | 3.35 | 0.47 | 0.66 | 0.29 | 0.20 | 0.02 |
| 151 | 1q0n_A | 1.22 | 0.79 | 0.62 | 0.73 | 0.28 | 0.27 | 0.01 |
| 160 | 1ako_A | 0.45 | 103.52 | 0.61 | 0.73 | 0.25 | 0.15 | 0.23 |
| 164 | 1ruv_A | 1.11 | 340.36 | 0.70 | 0.71 | 0.32 | 0.19 | 0.30 |
| 174 | 9pap_A | 0.13 | 15.44 | 0.28 | 0.52 | 0.23 | 0.04 | 0.10 |
| 216 | 1ca2_A | 1.30 | 16.69 | 0.51 | 0.64 | 0.22 | 0.27 | 0.08 |
| 252 | 1igs_A | 0.33 | 82.73 | 0.53 | 0.69 | 0.22 | 0.10 | 0.25 |
| 257 | 1xx2_A | 0.43 | 8.82 | 0.56 | 0.73 | 0.26 | 0.14 | 0.06 |
| 258 | 1sml_A | 1.81 | 1.22 | 0.71 | 0.76 | 0.50 | 0.25 | 0.01 |
| 290 | 1zio_A | 1.49 | 1.30 | 0.62 | 0.71 | 0.30 | 0.24 | 0.01 |
| 328 | 1lbm_A | 0.71 | 16.41 | 0.35 | 0.54 | 0.19 | 0.17 | 0.05 |
| 351 | 1snz_A | 0.65 | 183.94 | 0.46 | 0.61 | 0.23 | 0.19 | 0.18 |
| 362 | 1d6o_A | 0.32 | 80.27 | 0.69 | 0.76 | 0.27 | 0.07 | 0.38 |
| 376 | 1a4l_A | 0.09 | 31.44 | 0.20 | 0.43 | 0.23 | 0.03 | 0.16 |
| 394 | 1aj0_A | 1.68 | 28.26 | 0.66 | 0.74 | 0.24 | 0.35 | 0.07 |
| 444 | 1cel_A | 0.70 | 7.93 | 0.33 | 0.52 | 0.21 | 0.19 | 0.03 |
| 462 | 1pnt_A | 0.65 | 22.77 | 0.58 | 0.73 | 0.27 | 0.19 | 0.13 |
| 467 | 1cv2_A | 0.84 | 1.91 | 0.41 | 0.58 | 0.21 | 0.24 | 0.03 |
| 480 | 1czf_A | 0.72 | 16.61 | 0.40 | 0.57 | 0.19 | 0.17 | 0.07 |
| 597 | 1uch_A | 0.83 | 1.56 | 0.46 | 0.63 | 0.25 | 0.23 | 0.01 |
| 681 | 1gq8_A | 1.42 | 4.76 | 0.54 | 0.65 | 0.26 | 0.36 | 0.02 |
| 693 | 2rnf_A | 0.67 | 374.66 | 0.68 | 0.73 | 0.25 | 0.10 | 0.31 |
| 749 | 1nml_A | 0.96 | 1.37 | 0.29 | 0.48 | 0.31 | 0.20 | 0.01 |
| 814 | 1glo_A | 0.11 | 41.77 | 0.45 | 0.66 | 0.24 | 0.04 | 0.19 |
| 858 | 1mrq_A | 1.86 | 2.26 | 0.61 | 0.69 | 0.27 | 0.41 | 0.01 |
| 877 | 1pbg_A | 0.80 | 18.97 | 0.46 | 0.63 | 0.25 | 0.21 | 0.03 |
| 908 | 1rtu_A | 0.30 | 981.17 | 0.69 | 0.67 | 0.27 | 0.07 | 0.46 |
| 923 | 2acy_A | 1.21 | 43.56 | 0.42 | 0.57 | 0.21 | 0.21 | 0.11 |
| 931 | 2pth_A | 0.38 | 188.97 | 0.58 | 0.71 | 0.27 | 0.12 | 0.31 |

$a_S$: stability selection parameter; $a_A$: activity selection parameter; $D^2$: deviance explained by MSA model; $R$: Pearson correlation between observed and MSA-predicted lRMSD; sd($\phi_{\text{mut}}$), sd($\phi_{\text{stab}}$), sd($\phi_{\text{act}}$): standard deviations of mutation, stability, and activity SHAP components across residues
